# Transcriptome and interactome analyses identify the *TP53* interacting gene *RCCD1* as a candidate susceptibility gene at the 15p26.1 breast and ovarian cancer risk locus

**DOI:** 10.1101/2020.09.29.319699

**Authors:** Jasmine Plummer, Felipe Segato Dezem, Stephanie S. Chen, Subash Dhungana, Deepika Wali, Brian Davis, Justyna Kanska, Niko Safi, Ji-Heui Seo, Rosario I Corona, Joellen M Schildkraut, Paul DP Pharoah, Kate Lawrenson, Simon RV Knott, Matthew L Freedman, Siddhartha P Kar, Simon A Gayther

## Abstract

Common genetic variation in a region on chromosome 15q26 confers susceptibility to breast and ovarian cancer. The P53 interacting gene RCCD1 in this region is a candidate susceptibility gene for both cancers. In this study, a colocalization analysis of breast and ovarian cancer case-control genetic association studies in over 145,000 and 146,000 controls fine mapped the shared association in this region to 17 pleiotropic credible causal risk variants (*P*_breast_ < 1.16 × 10^−14^ and *P* _ovary_ < 7.50 × 10^−7^). These variants were strongly associated with the expression of RCCD1 in normal breast and ovarian tissues. Circular chromosome conformation capture (4C) analysis of *RCCD1* in breast and ovarian cancer cells identified similar patterns of *cis*-interaction and significant binding site enrichment for the *BRCA2* interacting gene *EMSY* (P_adjusted_ = 9.24 × 10^−6^). The 4C analysis pinpointed a single 2kB RCCD1 cis-interaction that contained two of the 17 shared risk variants. RCCD1 *trans*-interacting regions mapped to previously identified genome wide significant (P < 5 × 10^−8^) breast cancer risk loci (1p34.2 and 3p14.1) and to the pleiotropic breast-ovarian cancer risk locus at chromosome 9q34.2. Stable overexpression of RCCD1 in breast and ovarian cancer precursor cells identified 13 and 11 differentially expressed genes (DEGs) respectively associated with breast and ovarian cancer risk at genome-wide significance (*P*_MAGMA_ < 2.6 × 10^−6^ after Bonferroni correction). Eighty-two DEGs shared between breast and ovarian cancer were strongly enriched in TP53 (P = 9.9 × 10^−4^), Hippo (P = 2.51 × 10^−3^) and TNF signaling (P = 4.7 × 10^−3^) pathways.

## INTRODUCTION

Breast and ovarian cancers share common etiologies. Epidemiological studies have long established a role for sex steroid hormones (e.g. oral contraceptive use, hormone replacement therapy) influencing the risks of both cancer types^1,2^. Breast and ovarian cancers also share similar genetic liabilities suggesting a significant contribution of pleiotropy to risk of these cancers. In particular, the *BRCA1* and *BRCA2* genes are responsible for most families in which breast and ovarian cancers cluster together^3–7^. Pleiotropy is also an emerging feature of non-Mendelian forms of these and other cancers. Hundreds of common low penetrance susceptibility alleles have been identified for multiple cancer types using genome wide association studies (GWAS) and there is strong evidence that there is a shared genetic risk component for common alleles across cancers^8–11^. Common variants in several regions are associated with risks of both breast and ovarian cancer, suggesting that similar underlying biology may drive neoplastic development in both cancer types. For example, GWAS performed in breast and ovarian cancer populations have independently identified risk associations for each cancer at the 8q24.21 *CMYC* locus^12^, the 5p15.33 *TERT* locus^13^ and the 19p13.11 *ANKLE1/ABHD8* locus^14^. A recent meta-analysis combining GWAS data for both cancers identified shared (pleiotropic) risk regions that were not identified when each cancer was evaluated separately ^15^. This study performed a GWAS meta-analysis that included 62,533 women with breast cancer and 60,976 controls, and 15,437 women with invasive epithelial ovarian cancer and 30,845 controls and identified two new risk loci at 9q31.1 and 15q26.1 associated with both breast and ovarian cancer were at genome wide levels of significance in the combined breast and ovarian cancer data^15^.

The vast majority of disease associated variants lie in non-coding DNA regions and likely target tissue specific epigenomic elements that regulate the expression of a target susceptibility gene and/or critical gene expression networks involved in disease pathogenesis. Where the same genetic variants in a genomic region confer risk of two or more closely-related phenotypes, we hypothesize that there is shared underlying biology driving disease pathogenesis for the different phenotypes.

The goal of the current study was to interrogate the functional mechanisms underlying 15q26.1 breast and ovarian cancer risk locus and establish if the shared genetic associations identified for these cancers reflect a shared biology. Using the integrated approach illustrated in **Figure 1**, we applied a combination of genetic fine-scale mapping and epigenomic annotation, cell biology modeling of candidate genes identified by quantitative risk variant-gene expression assocation analysis, and *cis-* and *trans-*interaction studies in breast and ovarian cancer precursor and cancer cells to identify shared networks of differential gene expression and transcription factor binding linked to genetic risk.

**Figure 1:**
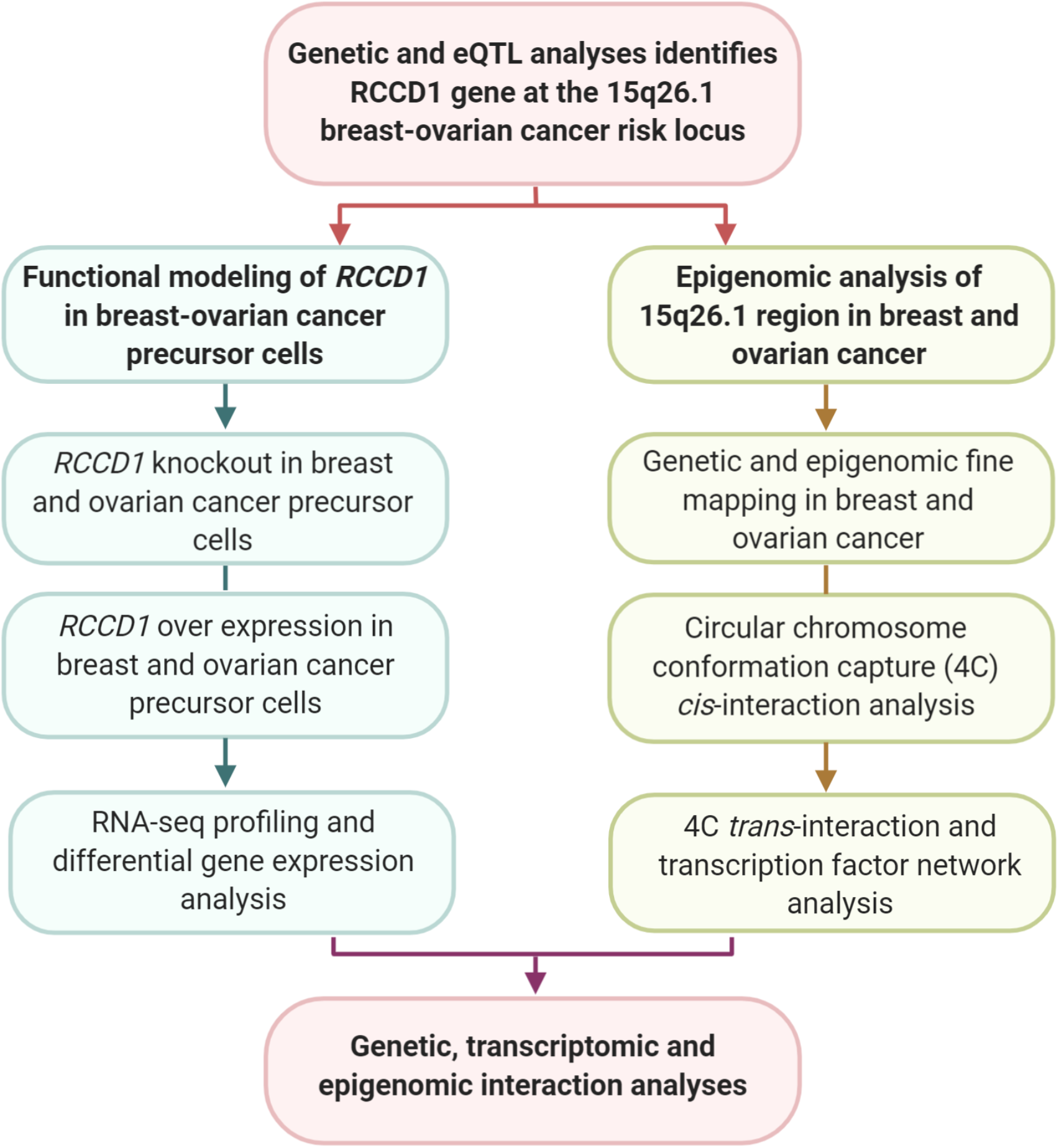
The experimental pipeline used for the genetic and functional analysis of the 15q26.1 (*RCCD1*) breast and ovarian cancer susceptibility locus.

## MATERIALS AND METHODS

### Genetic fine mapping analysis

SNP genotype and sample quality control, ancestry inference, imputation, genome-wide association and meta-analysis procedures for the breast and ovarian cancer GWAS meta-analysis data sets have been described previously^16,17^. All analyses were based on individuals of European ancestry and used 1000 Genomes Phase 3 (Version 5)-imputed or genotyped SNPs^18^. We harmonized effect and non-effect alleles and effect size estimate (beta coefficient) signs across the GWAS meta-analysis summary statistics data sets included in this study. We focused all analyses on 3,609 variants with minor allele frequency > 1% and imputation quality > 0.4 in each data set in the chr15:91009215-92009215 genomic region. Colocalization analysis was performed using the hypothesis prioritization in multi-trait colocalization (HyPrColoc) R package^19^. Expression Quantitative Trait Loci (eQTL) annotation of the credible set SNPs was conducted using the Qtlizer web application^20^ for the Genotype Tissue Expression (GTEx) project version 8 database ^21^ and using an online eQTL browser ^22^ for The Cancer Genome Atlas (TCGA)^23–25^.

### Cell culture

The FT282 and FT246 cell lines were established from fresh normal human fallopian tube (FT) tissue gifted by Dr. R. Drapkin^26^. These FT cell lines were maintained in DMEM-F12 medium (Gibco, Invitrogen, Carlsbad) supplemented with 10% fetal bovine serum (Gibco). Normal mammary epithelial cells (MCF10A, MCF12A) were cultured in MEBM medium supplemented with 100 ng/ml cholera toxin. Ovarian cancer cell line UWB 1.289 was purchased from ATCC and was cultured in a mixture (1:1) of 50% RPMI and 50% MEGM media with 3% FBS. The breast cancer cell line MCF7 was maintained in DMEM medium with 10% FBS. BT549 and Kuramochi were cultured in RPMI-1640 media supplemented with 10% FBS. All cell lines were thawed and allowed to recover for one day before expansion. All cell lines were passaged and plated onto 150mm plates. Cells were harvested at 80% confluency to be fixed, pelleted and snap frozen. Two biological replicates were included for each cell line.

### Preparation of 4C-Seq library

4C-Seq libraries were prepared using a protocol that has been modified from our previous method^27^. In brief, cells were single-cell suspended and their chromatin was crosslinked with 1% formaldehyde for 10 min at room temperature (21°C). Crosslinked cells were resuspended in a pre-chilled lysis buffer (10 mM Tris HCl, 10 mM NaCl, 0.2% NP-40) with fresh protease inhibitors (Millipore Sigma). The lysate was then digested with the primary restriction enzyme, DpnII (10,000 units/ml, New England Biolabs), and incubated overnight at 37°C. Next, DpnII-digested DNA was subjected to proximity ligation using 1X T4 DNA ligase (NEB), 1% TritonX-100, 0.1% BSA (NEB) and 4000U of T4 DNA ligase. Ligation was carried out overnight at 16°C followed by overnight cross-link removal with 20mg/ml Proteinase K (Qiagen). The samples were purified using phenol-chloroform and ethanol precipitated resulting in 3C libraries. The 3C libraries were subjected to a second restriction enzyme digestion using Csp6i (Life Technologies), followed by another proximity ligation using T4 DNA ligase. For each viewpoint, a total of 3.2 μg of the resulting 4C templates was used to carry out a scale-up inverse PCR, of which 16 reactions (200 ng in each) were amplified for 29 cycles and purified using Macherey Nagel Nucleospin Gel and PCR Purification kit (Takara Bio). Sequencing libraries were made of the 4C PCR products using Thruplex DNA-seq kit (Takara Bio). 4C libraries were subjected to Agencourt AMPure XP Bead cleanup (Beckman Coulter) using a bead-to-DNA ratio of 1:1 before high-throughput sequencing on the MiniSeq system (Illumina). Libraries were sequenced 1 × 75bp for a minimum coverage of 2 million read depth. Two technical replicates of each Dpn-Csp6I 3C library were made for each 4 cell lines.

### Primer design for 4C-Seq

The inverse primers were designed based on a viewpoint region. UCSC Genome Browser (build GRCh37/hg19) was used to locate the region of interest. Upon loading DpnII and Csp6I tracks, DpnII restriction sites flanking the region of interest were identified and the sequence between the nearest DpnII and Csp6I restriction sites were selected as the viewpoint region. Based on this region, 4C primers were designed with the following settings: optimal primer melting temperature of 57 °C (minimum 55 °C and maximum 60 °C); GC content between 40 and 60%. For DpnII-Csp6I libraries, 2 baits were designed as follows: 1) forward- TAGCTAGCACTGACAAGATC, reverse- GGATGTCCTCTGTTCTGAGT; and 2) forward- GAGGGCATAGGGTCAAGATC, reverse- CTTCAACCAGGCTCAGCT.

### 4C cis-analysis

For 4C raw sequencing data, cutadapt (version 2.0) was used to remove PCR primers (forward and reverse) and Illumina adapter from the reads, Burrows-Wheeler Aligner (bwa version 0.7.15-r1140) was employed to align raw sequencing files to the human genome (hg19). Samtools (version 1.9) was used to convert sequence alignment files (sam) to their binary version (bam). To determine statistically significant *cis*-interactions we used Bioconductor’s R package r3Cseq to count the reads per restriction fragment. Using technical replicates from each 4C library, the reads were normalized to RPM and interactions were obtained around 100kb from the view-point (4C bait), interactions closer than 3kb from the bait (promoter of RCCD1, coordinate 91496106 hg19) were removed. *Cis-*Interacting regions (CIRs) that were identified from biological replicates for each cell line, with a q-value < 0.01. Ovarian cancer CIRs were then merged based on the overlap of UWB1.289 and Kuramochi interactions. Breast CIRs were computed as the merged set of interactions in MCF7 and BT49 cell lines. Shared CIRs were identified by overlapping the breast and ovarian CIRs to obtain a unified *CIR* set with 100% of the interactions being shared across all four cell lines.

### 4C trans-analysis

For *trans-*interactions analysis raw coverage signal from sequencing files was inputted for FDR calculation, where a threshold of 0.01 was determined based on 100 permutations for each chromosome. From these permutation calculations, any window sizes that exceed 500 unique fragment ends were identified as *trans-* interactions^28^. All Interaction regions (*cis* and *trans*) were processed into bed files for plotting and visualization on UCSC Genome Browser as custom tracks.

### ChIP-Sequencing

Chromatin Immuno-Precipitation sequencing (ChIP-Seq) was performed as previously reported ^29^. Cells were fixed in 1% formaldehyde for 10 minutes and quenched with 2M glycine. Cells were harvested, lysed in buffer, and sonicated using the Covaris E220 evolution Focused-Ultrasonicator (Covaris). Antibodies used included H3K27ac (Diagenode), CTCF (Active Motif), H3K4me1 (abcam) and H3K4me3 (EMD Millipore). The H3k27ac antibody was incubated with 5 μg of chromatin while CTCF, H3k4me1 and H3k4me3 were incubated with 25ug of chromatin at 4°C overnight. Blocked magnetic Dynabeads (Life Technologies) were then added to the antibody-lysate conjugates and incubated at 4°C for 4 hours with rotation. Beads were then washed with RIPA buffer and treated with RNaseA and proteinase K (Qiagen). DNA was eluted from the beads in Tris-EDTA buffer and cleaned up using the Nucleospin Gel and PCR Purification kit (Macherey-Nagel). For every cell line, two independent immunoprecipitations and one input sample were submitted for next-generation sequencing on a HiSeq 5000 (Fulgent Genetics; Temple City, CA). All ChIP-Seq data was processed based on the ENCODE histone ChIP-Seq pipeline^30^ with modifications on the peak calling algorithm, to retain peaks with P< 1 × 10^−9^.

### Functional Annotation of Genome

The Statepaintr R package translates epigenomic files into chromatin segmentations using a rule-based-decision matrix combined with an abstraction layer with functional categories^31^. StatePaintR uses the deposited ChIP-Seq files of all uniquely mapping segment boundaries from the start and end coordinates of every peak then evaluates the presence or absence of each functional category. Next the package assesses overlaps of each segment to produce a boolean value. StatePaintR enables rank scoring of all states, allowing prioritization for non-coding variant annotation. ChIP-Seq data of the histone modifications (H3K27Ac-active enhancer/promoter), H3K4me3 (active promoter), H3K4me1(poised promoter/enhancer) and architectural complex protein (CTCF) was integrated for combinatorial patterning of overall chromatin states using Statepaintr in all precursor normal (MCF10A, FT246) and cancer cell lines (MCF7, BT549, UWB1.289 and Kuramochi). Bed files from these Statepaintr chromatin annotations were visualized on UCSC genome browser and IGV viewer.

### Transcription Factor Binding Site (TFBS) and protein interaction analysis

We utilized PERFECTOS-APE (PrEdict Regulatory Functional Effect of SNPs by Approximate P-value Estimation), a software built to identify TFs whose *in silico* binding sites are predicted to be significantly affected by a given nucleotide substitution^32^. This tool uses basic and dinucleotide Position Weight Matrices (PWMs) as TFBS and estimates the statistical significance (P value) of a predicted TFBS overlapping a SNP. Next, the method determines if the TFBS binding P values calculated for different homologous alleles differ significantly, such that extremely small or large ratios of P values are indicative of a TFBS existing for only one allele. TFBS motif variant prediction was conducted on the SNPs which intersected PCI/CCIs.

TFBS enrichment analysis was performed using ReMap 2020^33^, a curated database of 1135 transcription factors from 5798 ChIP-Seq experiments. To date this is the largest catalogue of experimentally derived regulatory regions by integrating all available DNA-binding assays generating 165M binding regions across the human genome. Genomic positions of the 4C interaction regions were crossed against the entire catalog of *in vitro* TF binding peaks. The collective chromosomal positions for all CCIs were compared to the ReMAP catalog of TF binding peaks to calculate statistical enrichment for TFs. First, *trans-*interacting regions were subdivided into 1kb peaks, in which these chromosomal positions were used as a query for the TFBS enrichment analysis. In this analysis a TF peak was considered for enrichment when the TF ChIP-Seq peak in the catalogue had 100% overlap within the 1kb query peak. The default setting for overlap is 10% but we selected 100% as the most stringent filter. For each queried peak, six genomic regions were shuffled to create a theoretical distribution. The p-value represents the probability to observe an effect at least as extreme as the result, under null hypothesis, which can be interpreted as an estimation of the false positive rate (FPR) and it is computed using the Poisson distribution (and validated empirically with randomized query regions). The q-value is the correction of the p-value to take into account the multiple testing due to the fact that the query is compared to each TF of the remap catalogue and it is an estimation of the false discovery rate (FDR), utilizing the Benjamini-Yekutieli method. Using ggplot2, the TFs enriched in CCIs were plotted by their Q Significance (-log10 of the Q-value) and frequency in which they occurred in the CCIs. This analysis was also applied to all PCIs to look for TF specifically enriched in these regions. TFs enriched in breast *trans*-interactions were evaluated using the union set of *trans*-interacting regions found in the breast cancer cell lines (MCF7 and BT549). Similarly the union set of *trans-*interactions in the ovarian cancer cell lines (UWB 1.289 and Kuramochi) were used as the regions queried in the ReMap dataset.

### Integrative modelling of chromatin landscapes and cistromes

We utilized Lisa (http://lisa.cistrome.org/) to predict the transcriptional regulators (TRs) of differentially expressed gene (DEG) sets. Briefly, Lisa uses publicly deposited histone mark ChIP-seq and chromatin accessibility profiles to construct a chromatin model related to the regulation of the DEG sets from breast and ovarian precursor models of RCCD1 overexpression. Using TR ChIP-seq peaks, this software analyses chromatin models to find the relevant TRs. to prioritize the candidate TRs, Lisa compares the predicted effects on the query and background gene sets using the one-sided Wilcoxon rank-sum test. We used two DEG sets as input for the model: set 1 was the DEG list from FT282^RCCD1+^(n= top 500; P<0,05; FDR<0.01) and set 2 was the DEG list in MCF12A^RCCD1+^ (n = 291; P<0,05; FDR<0.01). For the background gene set, we used all genes annotated on Gencode v.35. For the methods selection, ALL criterion was chosen which corresponds to both ISD-RP for motif and ChIP-Seq and TF ChIP-Seq Peak-RP.

### Luciferase reporter construct and activity assay

Reporter vectors were constructed in the pGL3-Promoter luciferase vector (Promega; GenBank Accession # U47298). The desired enhancer regions spanning the 4C interaction (chr15:91,504,893-91,507,207) were amplified from human gDNA by PCR using primer set F1/R3. pGL3-Promoter was linearized with XhoI restriction enzyme. Resulting two linear fragments was assembled using NEBuilder HiFi DNA Assembly kit (New England Biolab Inc.) as recommended by manufacturer to create pGL3-promoter-RCCD1 enhancer (pSD1) with an enhancer element containing risk alleles A for rs8028409 and T for rs8037137. pSD2, pSD3 and pSD4 containing one or both risk allele mutations (A to C and T to A for rs8028409 and rs8037137 respectively) were created using pSD1 as template to generate fragments for HiFi DNA assembly reaction introducing mutation at desired site/s. Primers used to generate fragments and introduce mutation/s are listed in **Supplementary Table S1**. Primer sets F1/R1, F2/R3, PGL3-F/PGL3-R generated 3 fragments for pSD2 plasmid assembly, primer sets F1/R2, F3/R3, PGL3-F/PGL3-R generated 3 fragments for pSD3 plasmid assembly and primer sets F1/R1, F2/R2, F3/R3, PGL3-F/PGL3-R generated 4 fragments for pSD4 plasmid assembly. All plasmid constructs were confirmed by Sanger sequencing. MCF7 and UWB1.289 cells were seeded in 6-well plates at a density of 0.3 million cells with respective media as mentioned above. Transfection was carried out when cells reached 60-70% confluency. 1.9µg firefly luciferase reporter plasmids (Promega; Accession # U47298) and 100ng of Renilla luciferase control plasmid (Promega; Accession # AF025848) was mixed with 3μL of BioT transfection reagent (Bioland Scientific LLC) and transfected the cell lines. Cells were grown in their corresponding media for 48 h. Transfected cells were assayed after 48 h (as indicated for firefly and Renilla luciferase activity using the Dual-Luciferase Reporter Assay System (Promega) in a GloMax® Explorer System (GM3500).

### RCCD1 overexpression plasmid construction and preparation

Overexpression of RCCD1 was accomplished using a sequence validated RCCD1 cDNA clone (HsCD00936904) This plasmid along with a control plasmid containing CAGG sequence were purchased from the DNASU repository (Biodesign Institute, Arizona State University, Tempe, AZ, USA). HEK 293T cells were plated onto 6cm plates and used to propagate the virus (RCCD1 overexpression plasmid and empty control plasmid). Twenty-four hours post plating, 30μl of BioT (Bioland Scientific, CA, USA) was added to 100 μL serum free media containing a total of 2μg per well of the desired RCCD1 expression construct or control empty vector plasmid. The BioT transfection and DNA solutions were then combined following manufacturer recommendations. Following 48 h of culture, virus was collected and prepared according to the manufacturer’s recommendations. Recipient cells (FT282 and MCF10A) were transduced with viral vectors. After 72h, transduction efficiency was evaluated and estimated by presence of fluorescence reporter from the plasmid.

### RNA isolation, library preparation and sequencing

Total RNA was isolated using RNA isolation kit (Machery Nagel) as per manufacturer’s instructions. RNA concentration and quality were determined by NanoDrop, Qubit and Agilent Bioanalyzer 2100. Total RNA were used as templates for cDNA libraries and were prepared as poly-A selected libraries using the Truseq stranded mRNA protocol (Illumina, San Diego, CA) and then sequenced on the Illumina Novaseq platform at AGCT Core (Los Angeles, CA) at 2 × 150 bp paired reads for a minimum of 35 million reads.

### RNA-sequencing analysis

Raw reads obtained from RNA-Seq were aligned to the transcriptome using STAR (version 2.5.0)^34^ / RSEM (version 1.2.25)^35^ with default parameters, using a custom human GRCh38 transcriptome reference downloaded from http://www.gencodegenes.org, containing all protein coding and long non-coding RNA genes based on human GENCODE version 23 annotation. All data analysis was performed using R and Bioconductor packages therein. For normalization purposes read count was transformed to TPM. Bioconductor’s package DESeq2 was used to compare the RCCD1 overexpression lines to control plasmid groups, performing differential expression analysis based on a model using the negative binomial distribution^36^.

### Cell proliferation assay

The human cell lines FT282^p53R175H^ (sgOR10A (Control ox), RCCD1 ox, sgRCCD1 (RCCD1 KO), CAGG ox (Control ox) and normal human breast epithelial cell MCF12A^p53R175H^ with CAGG ox, RCCD1 ox, RCCD1 KO and Control KO were grown in Dulbecco’s modified Eagle’s medium/Ham’s F12 medium supplemented with 10% FBS and Dulbecco’s modified Eagle’s medium and Ham’s F12 medium, 20 ng/ml Human epidermal growth factor, 100 ng/ml cholera toxin, 0.01 mg/ml bovine insulin and 500 ng/ml hydrocortisone, 95%; horse serum, 5% respectively. Cells were seeded in triplicate at the cell density of 10,000 cells/well in 96-well plate overnight in 200 ul respective media. After overnight incubation, equal volume of the 2X RealTime-Glo(tm) MT Cell Viability Reagent (Catalog # G9711)(Promega)) were added according to the manufacturer’s protocol. We measure the luminescence using the GloMax Microplate reader (Promega). To determine cell growth curve, we continue measuring luminescence at various times over the desired time course (e.g., 1hr, 2hr, 3hr, 4hr, 5hr, 8hr). We plot the luminescence versus different time points using linear regression curve fit. A one way ANOVA was used for statistical analysis of proliferation assays.

## RESULTS

### Genetic associations with breast and ovarian cancer risk in the 15q26.1 region colocalize to 17 candidate causal risk variants

We performed colocalization analysis of the previously identified breast and ovarian cancer susceptibility signal in the 15q26.1 region^15^ using summary genetic association data for 3,609 SNPs in the 1 Mb interval chr15:91009215-92009215 from the Breast Cancer Association Consortium (122,977 overall breast cancer cases and 105,974 controls)^16^ and the Ovarian Cancer Association Consortium (22,406 invasive epithelial ovarian cancer cases and 40,941 controls)^17^. Colocalization analysis supported two conclusions (**Figure 2A**): First, that there is a confirmed association with breast and ovarian cancer risk for common genetic variants in the chr15:91009215-92009215 interval (regional posterior probability of association (PPA) > 99%); second, that the alignment of cross-cancer associations in the region was strongly compatible (posterior probability of colocalization = 80%) with the presence of a single underlying causal signal that is associated with both breast and ovarian cancer risk.

**Figure 2:**
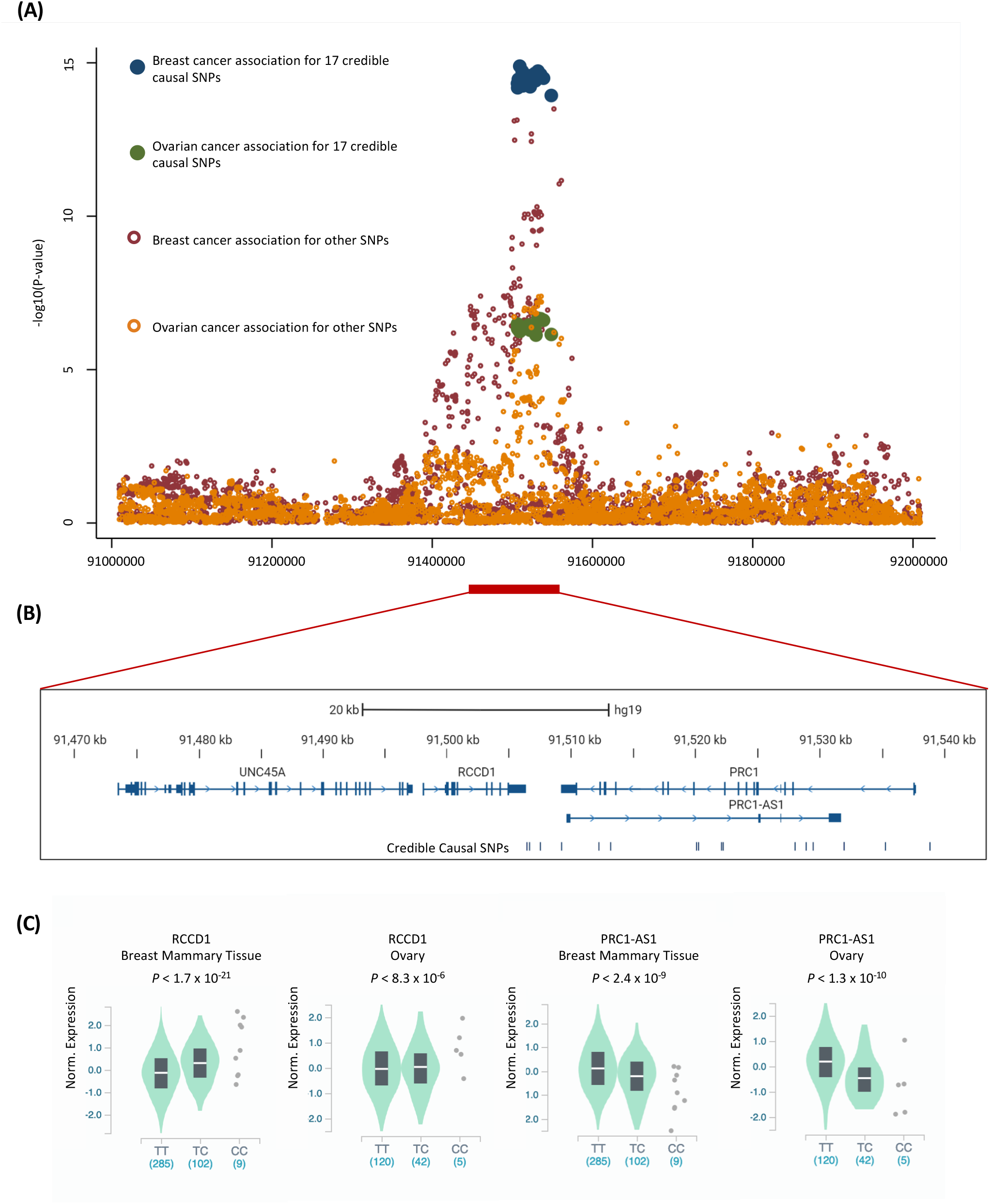
Fine mapping and expression quantitative trait locus analysis of the chromosome 15p26.1 breast and ovarian cancer risk locus: **(A)** Regional association plot for the chr15:91009215-92009215 interval derived from independent genetic association analysis performed breast and ovarian cancer case-control studies and from a meta-analysis that combines breast and ovarian cancer cases and controls to identify 17 credible causal risk variants that are shared between both cancers; **(B)** Regional localization plot showing the location of the 17 credible causal risk variants with respect to three nearest by genes, *RCCD1, PRC1* and the lncRNA *PRC1-AS1*; **(C)** Expression QTL analysis in normal breast and ovarian tissues from GTEx. Violin plots show allele-expression associations for rs8037137 index SNP. The T (risk associated) allele is significantly associated with decreased expression of *RCCD1* and increased expression of *PRC1-AS1* in both breast and ovarian tissues.

Colocalization analysis fine-mapped the pleiotropic breast and ovarian cancer signal in the region to a credible set of 17 SNPs that together explained > 99% of the shared association in the region (i.e., > 99% of the posterior probability of colocalization; **Figure 2B; Supplementary Table S2**). This joint cross-cancer fine mapping represents a 5 SNP (or 23%) improvement over a recently published large-scale, breast cancer-specific fine mapping effort that refined the breast cancer signal in the region down to 22 SNPs^37^. In a QTL based analysis of 396 breast and 167 ovarian tissues in the genotype-tissue expression (GTEx) project all 17 risk associated SNPs in the credible causal set were associated with the expression of two genes: *RCCD1* (*P*_breast_ < 1.7 × 10^−21^ and *P* _ovary_ < 8.3 × 10^−6^) and the long non-coding RNA *PRC1-AS1* (*P* _breast_ < 7.5 × 10^−9^ and *P* _ovary_ < 4.2 × 10^−9^) (**Supplementary Table S3**)^21^. The alleles associated with increased risk of breast and ovarian cancer were associated with decreased *RCCD1* expression and increased *PRC1-AS1* expression in both breast and ovarian tissues **(Figure 2C)**. The original lead SNP at this locus, rs8037137, was among the 17 SNPs in the credible causal set; gene expression associations for *RCCD1* and *PRC1-AS1* with rs8037137 genotypes are shown in **Figure 2C**. We also found that rs8028409 was the top eQTL for *RCCD1* expression in TCGA breast tumors (P = 3.2 x 10^−7^). As with our findings in GTEX, in TCGA data the A (risk) allele is associated with decreased RCCD1 expression. No expression associations were identified for these SNPs in ovarian tumors from TCGA^22^.

### RCCD1 interacts with shared epigenomic targets and two colocalized risk variants in breast and ovarian cancer cells

We hypothesized that *RCCD1* is the target susceptibility gene of genetic risk allele(s) at 15q26.1 regulated by the same genetic and epigenomic mechanisms in breast and ovarian cancers. To evaluate this, we first integrated the 17 risk variants with epigenomic features from profiling breast and ovarian precursor and cancer cell lines to identify putative regulatory targets of these variants **(Table 1)**. This identified intersections between 9/17 risk variants and different epigenomic features including active enhancers, poised enhancers, active regions and CTCF-bound regions, and with numerous transcription factors binding site (TFBS) motifs for 14/17 risk variants (**Table1; Supplementary Tables S4 and S5)**. We then used circular chromosome conformation capture (4C) analysis to identify which, if any, of these regions physically interacts with *RCCD1*. The promoter of *RCCD1* was used to bait interacting regions in the breast cancer cell lines MCF7 and BT549; and in the high grade serous ovarian cancer (HGSOC) cell lines UWB1.289 and Kuramochi. We identified a total of 65 *cis-*interacting regions (CIRs) with *RCCD1* including the same 9 CIRs in all four cell lines (**Figure 3A; Supplementary Table S6**). One of these nine CIRs in a region of approximately 2kb (chr15:91504892-91507207) harbored 2/17 credible causal risk variants: rs8028409 (*P*_breast risk_ = 4.73 × 10^−15^ and *P* _ovaran risk_ = 3.26 × 10^−7^) and rs8037137 (*P* _breast risk_ = 6.52 × 10^−15^ and *P* _ovarian risk_ = 4.61 × 10^−7^). Neither of these SNPs coincided with putative enhancers, promoters or CTCF-bound regions, but were predicted *in silico* to disrupt TFBS motifs for 16 factors (**Figure 3A; Table1; Supplementary Table S4**). We also used ReMap data to evaluate all local regions that interact with *RCCD1* for TFBS motif enrichment based on experimentally observed binding sites identified with ChIP-seq. CIRs were most significantly enriched for TFBSs with known roles in cancer including the BRCA2 interacting TF EMSY (P_adjusted_ = 9.24 × 10^−6^), the STAT1 and NF-κB regulator PIAS1 (P_adjusted_ = 6.74 × 10^−3^) and ZNF316 (P _adjusted_ = 6.74 × 10^−3^) **(Figure 3B)**.

**Table 1:** Details of the 17 credible causal risk variants (in order (5’ to 3’) of nucleotide location) identified by colocalization analyses of breast and ovarian cancer data at the 15q26.1 locus

| SNP ID | Nucleotide position (build 37) | P-Value breast cancer | P-Value ovarian cancer | SNP Score <sup>a</sup> | # TF motifs intersections | Tissue specific epigenomic intersections <sup>b</sup> |
| --- | --- | --- | --- | --- | --- | --- |
| *rs8028409* <sup>c</sup> | **91506422** | **4.73 x 10<sup>-15</sup>** | **3.26 x 10<sup>-7</sup>** | **0.046** | **13** | - |
| *rs8037137* <sup>c</sup> | **91506637** | **6.52 x 10<sup>-15</sup>** | **4.61 x 10<sup>-7</sup>** | **0.023** | **3** | - |
| rs112770790 | 91507495 | 3.44 x 10<sup>-15</sup> | 4.97 x 10<sup>-7</sup> | 0.039 | 7 | EPR |
| rs77554484 | 91509215 | 1.28 x 10<sup>-15</sup> | 5.73 x 10<sup>-7</sup> | 0.090 | - | AR |
| rs2290202 | 91512267 | 1.87 x 10<sup>-15</sup> | 4.38 x 10<sup>-7</sup> | 0.073 | 11 | - |
| rs2890156 | 91513157 | 5.65 x 10<sup>-15</sup> | 4.81 x 10<sup>-7</sup> | 0.022 | 14 | - |
| rs12595025 | 91520131 | 4.03 x 10<sup>-15</sup> | 4.20 x 10<sup>-7</sup> | 0.039 | 15 | - |
| rs12594925 | 91520287 | 2.64 x 10<sup>-15</sup> | 3.54 x 10<sup>-7</sup> | 0.070 | 16 | CTCF |
| rs113343095 | 91522092 | 3.37 x 10<sup>-15</sup> | 4.54 x 10<sup>-7</sup> | 0.056 | 16 | AR; EAR |
| rs8026714 | 91522253 | 6.05 x 10<sup>-15</sup> | 5.20 x 10<sup>-7</sup> | 0.021 | - | EPR; AR; EAR |
| **rs2301825<sup>d</sup>** | **91528070** | **2.24 x 10<sup>-15</sup>** | **4.37 x 10<sup>-7</sup>** | **0.070** | **4** | **EAR** |
| rs150333865 | 91528875 | 3.83 x 10<sup>-15</sup> | 6.46 x 10<sup>-7</sup> | 0.026 | 24 | EAR |
| rs60290423 | 91529452 | 3.21 x 10<sup>-15</sup> | 7.50 x 10<sup>-7</sup> | 0.027 | 4 | EPR |
| rs12594752 | 91531995 | 1.86 x 10<sup>-15</sup> | 2.13 x 10<sup>-7</sup> | 0.159 | 1 | - |
| rs76119208 | 91535329 | 2.40 x 10<sup>-15</sup> | 2.20 x 10<sup>-7</sup> | 0.129 | 29 | EPR |
| rs78029804 | 91538920 | 3.17 x 10<sup>-15</sup> | 2.46 x 10<sup>-7</sup> | 0.098 | - | - |
| rs3826033 | 91548601 | 1.16 x 10<sup>-14</sup> | 7.14 x 10<sup>-7</sup> | 0.008 | 15 | - |
| <sup>a</sup> Ranking score for the top 17 risk associated SNPs in breast and ovarian cancer from co-localization analysis (i.e., posterior probability of colocalization at each SNP). These 17 SNPs explain > 99% of the shared association in the region<br><sup>b</sup> EAR: Active enhancer region; EPR: Poised enhancer region; CTCF: Architectural complex region; AR: Active region<br><sup>c</sup> Intersects a 4C interacting region in breast and ovarian cancer cells (Figure 3A)<br><sup>d</sup> Intersects a 4C interacting region in breast and ovarian precursor cells (Figure 3A) |  |  |  |  |  |  |

**Figure 3:**
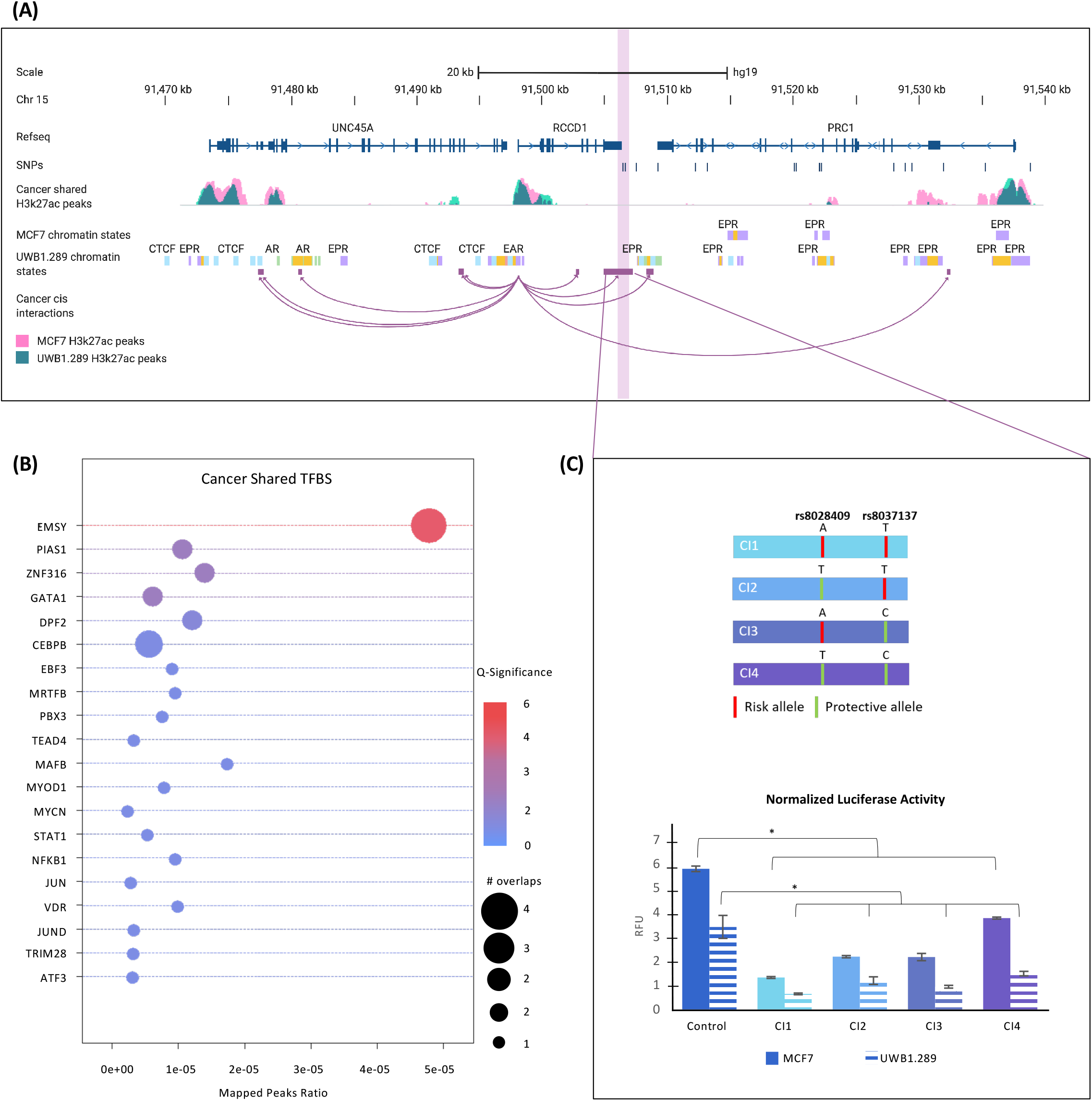
Circular chromosome conformation capture (4C) analysis anchored to *RCCD1* in breast and ovarian precursor and cancer cell lines. **(A)** Genomic locus plot of chr 15: 91470-91450 showing. RefSeq genes annotated from hg19 build of UCSC genome browser. Shared 4C interactions with RCCD1 in breast and ovarian cancer cell lines (magenta loops) and breast and ovarian precursors (blue loops). Combined tracks of enhancer-associated H3K27ac histone marks catalogued in breast (pink peaks) and ovarian (dark teal peaks) cancer and precursor cell lines show the 4C-RCCD1 landscape extending upstream and downstream of RCCD1, with respect to the colocalized credible causal risk variants (vertical dark blue lines).Interacting peaks that overlap 2/17 risk variants (rs8037137 and rs8028409) in cancer cell lines and 1/17 risk variants (rs2301825) in precursor cell lines are highlighted by the pink and purple vertical bars respectively. Chromatin states annotated in breast (MCF7) and ovarian (UWB1.289) cancer cell lines: Active region (AR): active promoter region (PAR); poised promoter region (PPR); active enhancer region (EAR); poised promoter region (PPR); architectural complex region (CTCF). **(B)** Enrichment of TFs ChIP-Seq peaks found in *cis*-interactions of breast cancer (MCF7, BT549) cell lines. Color designates the statistical significance (Q value) of the TF enrichment with highest (red) to lowest E value (blue). Circle sizes depict the number of overlaps the TFs intersect with *cis*-interactions; **(C)** Enrichment of TFs ChIP-Seq peaks found in *cis*-interactions of (B) ovarian cancer (UWB1.289, Kuramochi) cell lines.

We tested the functional effects of different allelic combinations of rs8028409 and rs8037137 on RCCD1 expression using luciferase reporter assays in breast (MCF7) and ovarian (UWB1.289) cancer cells. Four constructs each containing different allelic combinations of rs8028409 (A-risk/T-protective) and rs8037137 (T-risk/C-protective) were transfected into MCF7 and UWB1.289 cells. Constructs containing only the risk alleles of rs8028409 and rs8037137 were associated with the lowest RCCD1 expression in both breast and ovarian cancer cells. Constructs expressing only protective alleles were associated with the highest RCCD1 expression (*P*_breast_ <0.0001 and *P*_ovarian_ = 0.02 respectively) while constructs containing different combinations of protective/risk allele for each SNP were associated with intermediate RCCD1 expression **(Figure 3C)**.

### RCCD1 4C trans interactions in breast and ovarian cancer risk are enriched for disease associated transcription factor binding sites

We used 4C data in the breast and ovarian cancer cells to identify long-range *trans* interacting regions (TIRs) between RCCD1 and regulatory elements throughout the genome. In total, we found 165 TIRs across all four cancer cell lines with 30 and 46 TIs respectively identified in MCF7 and BT549 breast cancer cells, 66 and 23 TIRs respectively identified in UWB1.289 and Kuramochi ovarian cancer cells and three TIRs shared across all four cell lines (**Supplementary Table S7)**. Five other TIRs were shared between both breast cancer cell lines, and nine other TIRs were shared between both ovarian cancer cell lines **(Figure 4A)**. We performed enrichment analyses of these TIRs with respect to TFBS motifs from ReMap data. There were significant enrichments for many TFBSs shown to play a role in cancer development generally and breast and ovarian cancer biology specifically. For breast cancer TIRs, the most significantly enriched TFBS motifs included the transcriptional regulator chromobox protein (CBX1), the P53 degradation E3 ligase protein (RNF2), the DNA double-strand break repair protein RAD21 and the retinoblastoma protein (RB1)**(Figure 4C; Supplementary Table S8)**. For ovarian cancer, the most significantly enriched TFBS motifs included the MYC Associated Factor X (MAX) protein, the stromal antigen 1 protein(STAG1) and bromodomain transcriptional regulatory (BRD4) **(Figure 4D; Supplementary Table S9)**.

**Figure 4:**
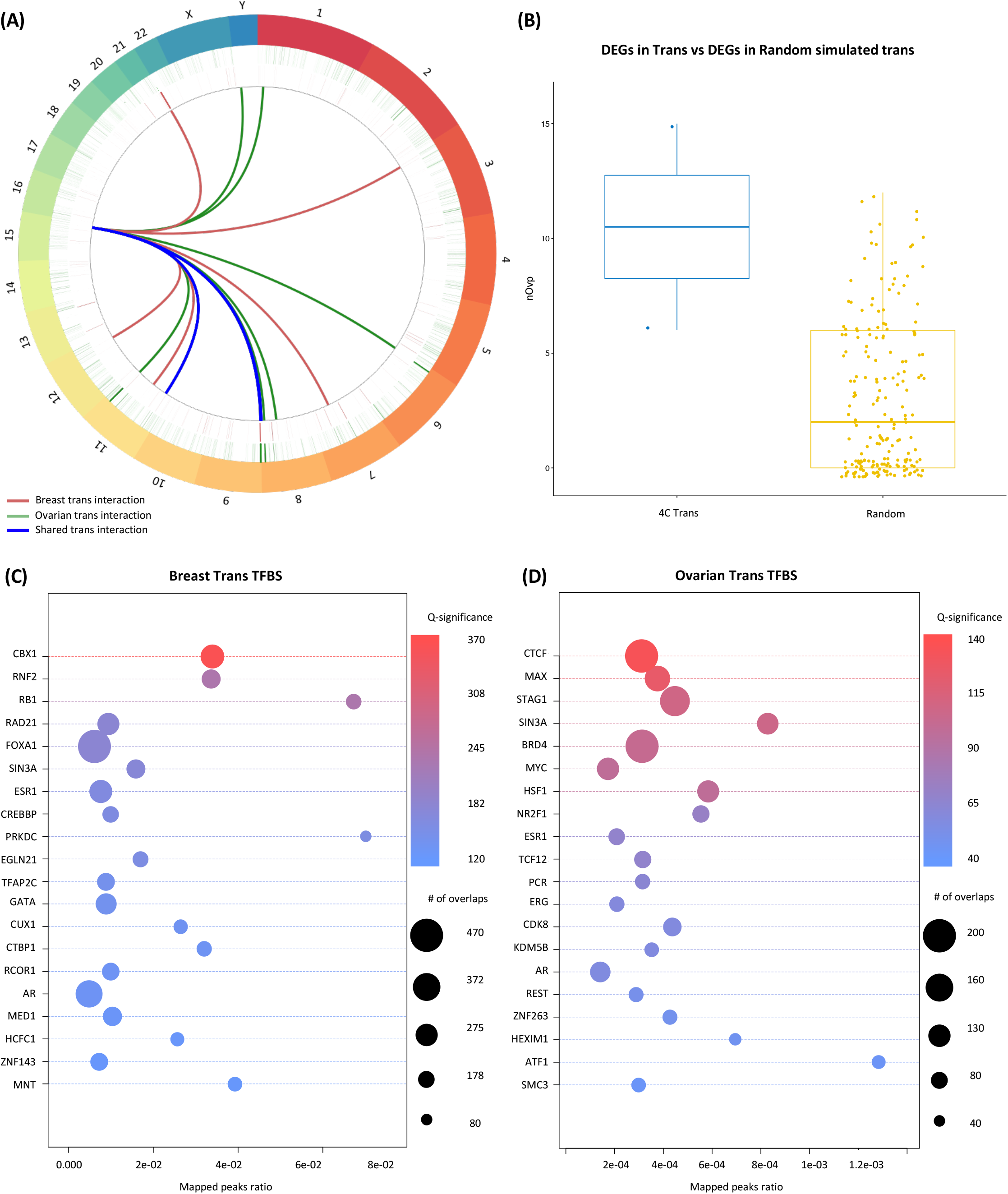
*Trans*-4C interaction and network analysis for RCCD1 in breast and ovarian cancer cells. **(A)** Circos plot of genome wide RCCD1 *trans*-interactions (FDR<0.01) in breast cancer cell lines (MCF7, BT549) and ovarian cancer cell lines (UWB.1289, Kuramochi) are depicted as red and green lines, respectively, looping in the center. Interactions are aligned with respect to chromosomes (outer track), and breast and ovarian cancer GWAS risk loci (P≥5×10^−8^) which intersect *trans*-interactions (inner circle). **(B)** Wilcoxon text of DEGs enriched in RCCD1-TIRs in breast and ovarian cancer cells based on the simulation of more than a thousand random permutations (P = 0.049)**(C)** Enrichment of TF ChIP-Seq peaks found in *trans*-interactions of breast cancer cell lines MCF7 and BT549. Color of the circles represent the statistical significance of the TF enrichment with red representing the highest and blue the lowest. Circle sizes represent the total numbers of overlaps between TFs and *trans*-interactions in breast cancer; **(D)** Enrichment of TF ChIP-Seq peaks found in *trans*-interactions of ovarian cancer cell lines UWB1.289 and Kuramochi. Color codes are the same as in (B).

### Phenotypic and molecular analysis in breast and ovarian cancer precursor cells support a role for RCCD1 in early stage neoplastic development

Expression QTL analyses suggest that the risk alleles of variants at 15q26.1 are associated with a decrease in *RCCD1* expression (**Figure 2C**); but in breast and ovarian tumors and in cancer cell lines the evidence suggests that *RCCD1* is overexpressed during tumor development (**Supplementary Figure 1A**)^23,24^. Therefore, we tested the effects of modulating *RCCD1* expression by both overexpressing and knocking down the gene separately in models of breast (MCF12A) and ovarian (FT282) cancer precursor cells. We first generated single cell clonal models of *RCCD1* overexpression (MCF12A^RCCD1+^ and FT282^RCCD1+^) and confirmed overexpression of both *RCCD1* transcript and protein (**Supplementary Figure 1B**). MCF12A^RCCD1+^ and FT282^RCCD1+^ cells were significantly more proliferative than MCF12A and FT282 cells (P = 1.7 × 10^−3^and 5.1 × 10^−3^ respectively) (**Figure 5A & B**). Next, we used CRISPR/Cas9 to knockdown *RCCD1* expression (MCF12A^RCCD1-^ and FT282^RCCD1-^), derived single cell clones and confirmed loss of *RCCD1* transcript and protein expression (**Supplementary Figure 1C**). *RCCD1* knockdown induced a strong growth suppression in MCF12A^RCCD1-^ cells (P = 4.5 × 10^−3^ ; **Figure 5A**) but an increase in proliferation in FT282^RCCD1-^ cells (P = 1.1 × 10^−3^; **Figure 5B**).

**Figure 5:**
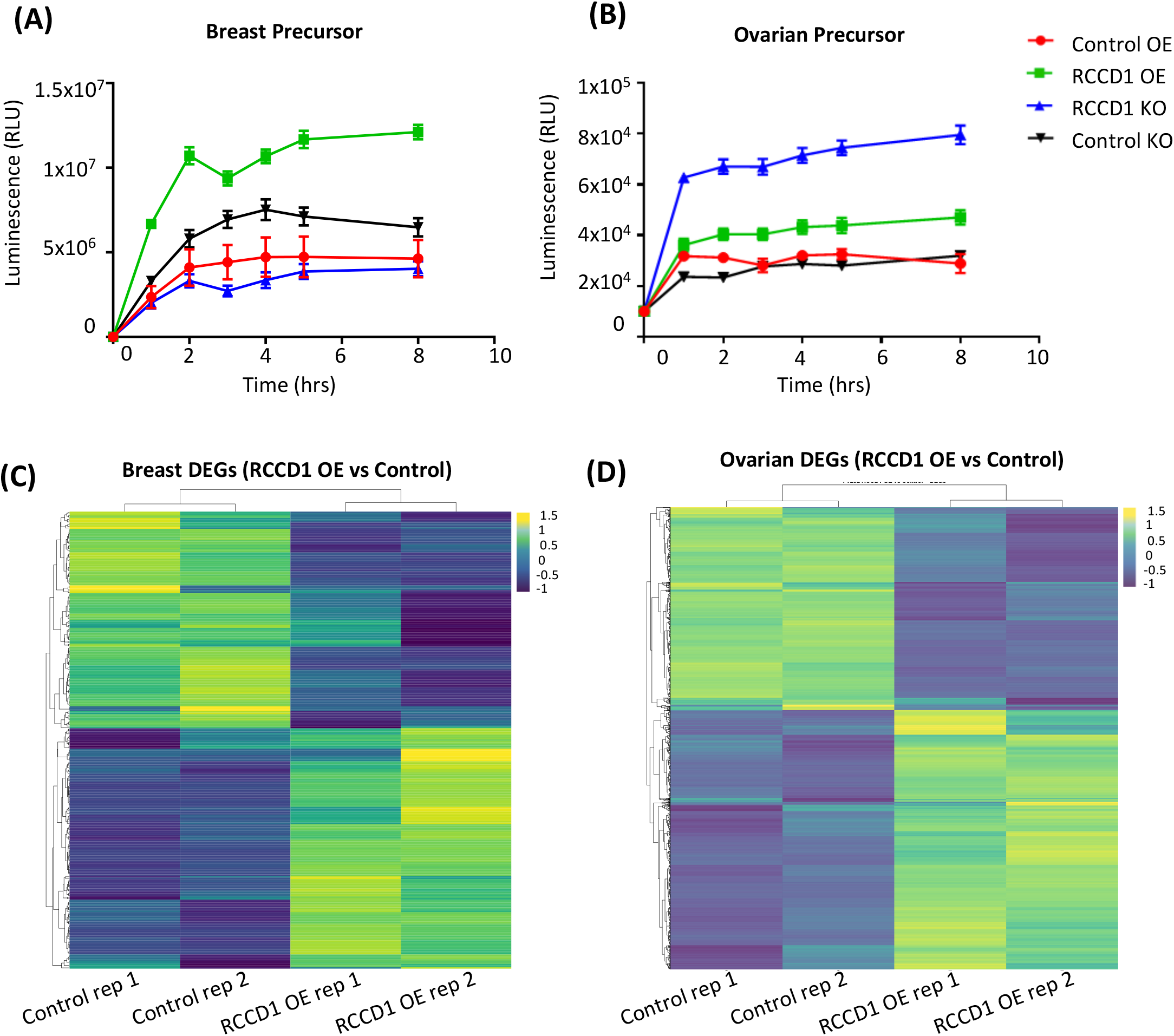
Phenotypic analysis of RCCD1 overexpression models in precursor breast and ovarian cancer cells. **(A)** Cell proliferation is significantly increased in precursor (**A)** breast (MCF12A RCCD1 OE) and (**B)** ovarian (FT282-RCCD1 OE) cell models of RCCD1 overexpression compared to control (CONTROL OE). Cell proliferation decreased in RCCD1 knockout cell models in breast (MCF12A RCCD1 KO) but increased in ovarian cancer models (FT282 RCCD1 KO) compared to control cells (OR104A KO). Heat map of color coded expression of DEGs (| log2FC |> 0.5, p_adjusted_<0.05) from overexpression of RCCD1 in **(C)** breast cancer precursor cells (MCF12A^RCCD1+^) and **(D)** ovarian cancer precursor cells (FT282^RCCD1+^). Upregulated genes are depicted in yellow and down regulated genes are shown in green.

Because the phenotypic effects associated with RCCD1 overexpression were similar in MCF12A and FT282 cells, we chose to perform whole transcriptome profiling of MCF12A^RCCD1+^ and FT282^RCCD1+^ clones by RNA-seq. We identified 291 differentially expressed genes (DEGs) (|log2FC | >0.5, P _adjusted_< 0.05) between MCF12A and MCF12A^RCCD1+^ cells of which 138 genes were up-regulated and 153 genes were down-regulated **(Figure 5C, Supplementary Table S10)**. Comparing FT282 with FT282^RCCD1+^ cells, we identified 2,469 DEGs (| log2FC |> 0.5, p_adjusted_< 0.05), of which 1,084 genes were up-regulated and 1,385 genes were down-regulated in **(Figure 5D, Supplementary Table S10)**. Eighty-two DEGs were shared between breast and ovarian cancer precursor cell line models. All DEGs were annotated using the KEGG database to identify biologically relevant pathways (**Supplementary Table S11-13)**. In MCF12A^RCCD1+^ models we identified significant enrichments for pathways involved in PPAR (peroxisome proliferator-activated receptor) signaling (P = 0.01), the pro-inflammatory cytokine IL-17 (interleukin 17A) signaling (P = 0.02) and steroid hormone biosynthesis (P = 0.03) **(Supplementary Table S11)**. In FT282^RCCD1+^ cells we identified significant enrichments for pathways involved in Ras (P = 9.4 × 10^−5^), Rap1 (P = 9.83 × 10^−5^) and p53 signaling pathways (P = 9.9 × 10^−5^) **(Supplementary Table S12)**. Pathway analyses of the 82 DEGs that were shared between breast and ovarian *RCCD1* overexpression models identified significant enrichment of genes in the TP53 (P = 9.9 × 10^−4^), Hippo (P = 2.51 × 10^−3^) and TNF signaling (P = 4.65 × 10^−3^) pathways **(Supplementary Table S13)**.

### Evidence that RCCD1 interacts with and operates in the TP53 pathway in breast and ovarian cancer

We further explored the putative role of RCCD1 in the TP53 pathway. RCCD1 is a known partner of the histone H3K36 demethylase KDM8 (JMJD5)^38^ which interacts with the DNA-binding domain of TP53 to negatively regulate its activity (**Figure 6A**). This finding was mainly driven by enrichment for TP53-associated DEGs identified in FT282^RCCD1+^ models. Several DEGs identified in these models are closely related partners of TP53 signaling including: MDM2, a critical negative regulator of P53; the cyclin-dependent kinase inhibitor CDKN2A; and CCND1 and CCND2, two regulators of cyclin-dependent kinases (**Figure 6B**). In two independent clones of RCCD1 knockdown in FT282 cells stably expressing a functional mutation of TP53 (R175H), mutant TP53 was significantly depleted (**Figure 6C**). We analyzed RCCD1 expression with respect to somatic, functional TP53 mutations in primary breast and ovarian cancers using TCGA data^39^. Both breast and ovarian cancers with increased RCCD1 expression were significantly more likely to harbor functional TP53 mutations than tumors with low RCCD1 expression (*P* _breast_ = 8 × 10^−12^ and *P* _ovary_ = 4 × 10^−3^) (**Figure 6D**).

**Figure 6:**
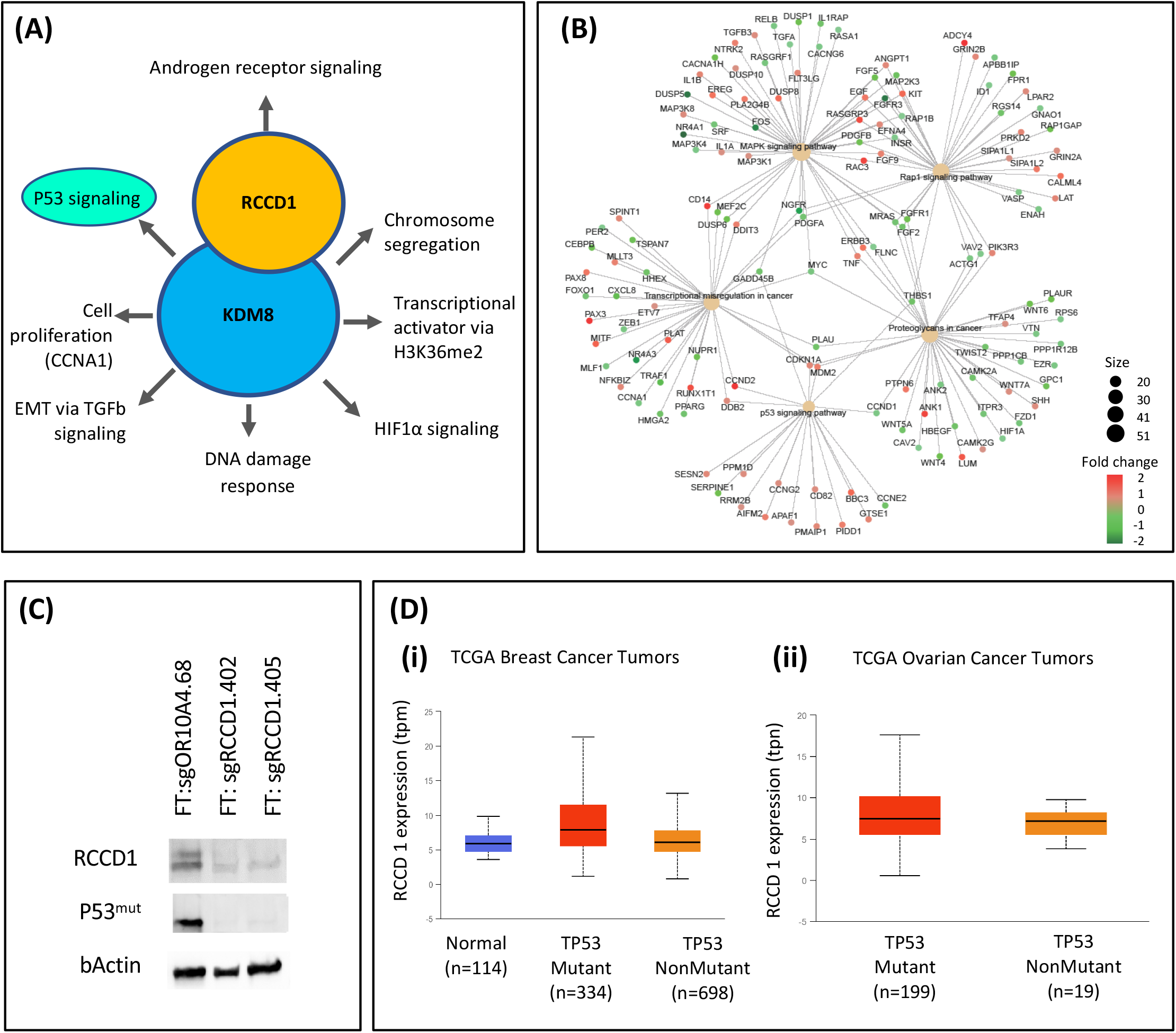
RCCD1 is a putative regulator of the P53 pathway in breast and ovarian cancer cells. **(A)** RCCD1 operates in the P53 pathways through its interaction with KDM8. As part of this complex RCCD1 also regulates HIF1A signaling, transcriptional activation via H3K36me2, epithelial mesenchymal transition via TGF beta signaling, cell proliferation via CCNA1, DNA damage response and Androgen receptor signaling. **(B)** Pathway analysis of DEGs from RCCD1 overexpression models are enriched in key proteins of the TP53 signaling pathway. **(C)** Western blot analysis after CRISPR/Cas9 knockout of RCCD1 in fallopian tube epithelial cells expressing the R175H P53 mutation (FT282^P53:R175H^). Two different RCCD1 knockout clones are compared to knockdown of a control gene, the olfactory receptor OR10A4. After RCCD1 knockdown and clonal selection, mutant TP53 is depleted in FT282^P53:R175H^ cells indicating downregulated expression of mutant TP53 is selected for in the presence of RCCD1 knockout. **(C)** RCCD1 expression in TCGA breast and ovarian tumors based on TP53 mutation status. These plots were obtained using the UALCAN web tool (http://ualcan.path.uab.edu/cgi-bin/TCGAExResultNew2.pl?genenam=RCCD1&ctype=BRCA. Briefly, in UALCAN, TP53 mutation status was obtained from TCGA whole exome sequencing data. Mutation Annotation Format files (derived from VarScan2) from the Genomic Data Commons portal were matched with TCGA RNA-seq data and the analysis performed stratifying the samples with/without TP53 mutation.

### Differentially expressed genes identified in RCCD1 overexpression models are associated with breast and ovarian risk alleles

We used GWAS data generated by the Breast and Ovarian Cancer Association Consortia to map common variants across the genome to genes and performed gene-level association analysis using MAGMA^40,41^. This identified 281 genes associated with breast cancer risk and 42 genes associated with ovarian cancer risk at genome-wide significance (*P*_MAGMA_ < 2.62 × 10^−6^; threshold adjusted for testing 19,100 genes). Thirteen of the 281 breast cancer associated genes and 11/42 ovarian cancer associated genes from MAGMA showed significant gene expression perturbations (P_adjusted_ < 0.05) in MCF12A^RCCD1+^ cells and FT282^RCCD1+^ cells, respectively (counts exclude *RCCD1*; **Supplementary Table S14**). There was a |log2FC| > 0.5 fold change in expression of *TRIM31* in the MHC region and *CDYL2* and *NOTCH2NL* in MCF12A^RCCD1+^ cells and *ANKLE1, BNC2, HOXB3*, and *LRRC37A* in FT282^RCCD1+^ cells. Several differentially expressed genes in FT282^RCCD1+^ cells were located at pleiotropic breast and ovarian cancer risk loci, such as *ANKLE1* and *ANO8* at the 19p13.11 locus^14^; *LRRC37A, KANSL1, ARL17A*, and *PLEKHM1* at the 17q21.31 locus^42,43^, and *TERT* at the 5p15.33 locus ^13^. The *HOXB* cluster and *BNC2* have previously been identified as having specific functional roles in mediating ovarian cancer risk^44,45^. *MLLT10* was differentially expressed in both MCF12A^RCCD1+^ and FT282^RCCD1+^ cells (P _adjusted_ < 0.05) and was significantly associated with both breast and ovarian cancer risk in the MAGMA analyses (*P* _MAGMA_ < 2.62 × 10^−6^).

### Integrating transcriptomic and RCCD1 interactome data validates 4C trans-interactions identified in breast and ovarian cancers

To validate the functional significance of RCCD1-TIRs in breast and ovarian cancers, we integrated these regions with the differential expressed genes identified after overexpression of *RCCD1* in MCF12A and FT282 cells. We identified six DEGs(PCBP3, BSCL2, COL18A1, NUP210, POLR2G and PARP10) in MCF12A^RCCD1+^ cells that overlap TIRs in breast cancer cell lines and eighteen DEGs (EEF1G, BCDIN3D, TUBD1,PPM1D, RNF43, BCAS4, TSHZ2, DOK5 DOCK3, MKRN2OS, HRH1, RPL26L1, VDAC1, SKP1, ZDHHC11, RPS10, PRKRIP1, RASA4) that overlap TIRs in ovarian cancer cell lines. This enrichment of DEGs in RCCD1-TIRs in breast and ovarian cancer cells was significantly more than expected by chance based on the simulation of more than a thousand random permutations (Wilcoxon P = 0.049; **Figure 4B)**.

Finally, we used the online tool LISA (Cistrome) to evaluate the TFs and chromatin regulators that may be responsible for the perturbation of DEGs found in MCF12A^RCCD1+^ and FT282^RCCD1+^ cells. In MCF12A^RCCD1+^ cells 13 out of the top 50 TFs overlapped ReMAP and Cistrome predictions (including YY1, HDAC2, EGR1, STAG1, HEXIM1, MBD3, ELF1 and CHD2) **(Supplementary Table S15)** and in FT282^RCCD1+^ cells 11 of the top 50 TFs overlapped ReMAP and Cistrome predictions from the DEGs identified in FT282^RCCD1+^ cells (including NR2F1, SP1, ERG, ESR1, RUNX1 and HIF1A) **(Supplementary Table S16)**. We also identified several TFs that were shared between breast and ovarian cancer cells in both our trans TF enrichment and Cistrome analyses including MYC, RAD21, CTCF, MAX and AR.

## DISCUSSION

We have shown that the genetic architecture of common variant risk associations in the 15q26.1 region is the same in both breast and ovarian cancer suggesting that similar functional mechanisms at this locus drive disease development for both cancer types. We focused on RCCD1 (regulator of chromosome condensation domain-containing protein 1) in this region as the most likely breast and ovarian cancer susceptibility gene and target of risk variation following expression quantitative trait locus (eQTL) analysis which found significant associations between risk genotypes and *RCCD1* expression in normal breast and ovarian tissues. However, we cannot rule out that other genes in the region may be the target susceptibility gene operating in collaboration with or independently from *RCCD1* and our eQTL analyses also identified significant associations with the lncRNA *PRC1-AS1* (Protein Regulator of cytokinesis 1- antisense 1) in normal breast and ovarian tissues. *PRC1-AS1* overlaps the *PRC1* gene, and as such may play a role in regulating *PRC1* transcript expression. While we did not identify a significant eQTL for *PRC1* in this analysis, a previous transcriptome wide association study (TWAS) has identified a significant splice-QTL for *PRC1* in breast tumors which fully explained the GWAS signal in ovarian cancer at the 15q26.1^46^. In addition, recent studies have suggested that *PRC1* overexpression may be associated with the development of multiple tumor types^47–49^ and in one study *PRC1* mRNA and protein expression were upregulated in high-grade serous ovarian cancers, particularly in tumors from patients without *BRCA1/BRCA2* mutations^50^.

*RCCD1* is a highly plausible breast and ovarian cancer susceptibility gene. By partnering with the histone H3K36 demethylase KDM8 (JMJD5)^38^,RCCD1 interacts with the DNA-binding domain of TP53 to negatively regulate its activity, and is an important complex for spindle organization, chromosome segregation and accurate mitotic division ^38^. Germline pathogenic mutations in TP53 are responsible for the autosomal dominant Li-Fraumeni syndrome (LFS), which is most commonly associated with early onset sarcoma, and malignancies of the brain, breast and adrenal glands^51^. The risk of breast cancer in women with LFS is estimated to be about 50% by age 60^52^. Somatic P53 mutations are also essential for HGSOC development, are common aberrations in triple negative breast cancer^25,24^, and are associated with *BRCA1* and *BRCA2* mutant cancers^53,54^. In this study, we provide functional evidence that RCCD1 interacts with TP53 in early stage neoplastic development. Knockout of wild-type *RCCD1* expression in fallopian tube cells expressing the functional *TP53* hotspot mutation (R175H) resulted in depletion of TP53 protein expression. Consistent with this, primary breast and ovarian tumors analyzed by TCGA ^39^ with low *RCCD1* expression are significantly less likely to also harbor pathogenic TP53 mutations than tumors with higher *RCCD1* expression.

The interaction with TP53 suggests that *RCCD1* plays an important role in cell cycle regulation and DNA damage repair^55^. Other data from this study support this hypothesis. The same *RCCD1 cis*-interacting regions identified in both breast and ovarian cancer cell lines were significantly enriched for *EMSY* binding site motifs. *EMSY* is a transcriptional repressor that directly interacts with the DNA double strand break repair protein BRCA2, a well characterized breast and ovarian cancer susceptibility gene. Thus, EMSY likely dysregulates DNA damage repair to induce genomic instability. Immunofluorescence analysis in the normal mammary epithelial cell line MCF-10A and breast cancer cell line MCF-7 shows that EMSY localizes to the nucleus after exposure to ionizing radiation; there is also direct evidence that EMSY locates to sites of DNA lesions after DNA damage^56,57^. In another study, overexpression of *EMSY* disrupted the BRCA2/RAD51 pathway in response to DNA-damage, leading to the hypothesis that amplification of EMSY mimics BRCA2 deficiency by overriding RPA and PALB2, crippling the BRCA2/RAD51 complex at site of DNA damage^58^.

*RCCD1 trans*-4C interaction analyses identified genomic regions that were highly significantly enriched in motifs for TFs that are known to be involved in breast and/or ovarian cancer biology including in the three shared regions identified in four breast and ovarian cancer cell lines. This included *RAD21* an essential gene that functions in the DNA double strand break repair pathway, further supporting the putative functional role of *RCCD1* in DNA repair. A recent study has shown that mutant p53 stimulates cell invasion through its interaction with RAD21 in ovarian cancer cells, while *RAD21* expression is associated with poor prognosis in *BRCA2* associated breast cancers ^59^. ESR1 (estrogen receptor 1) was also enriched in *RCCD1 trans*-interacting regions in breast and ovarian cancer cells. *ESR1* is a nuclear receptor that is activated by the sex hormone estrogen. *ESR1* aberrations play a significant role in the development of breast cancers and their progression to metastasis and resistance^60,61^. Germline variants at the *ESR1* locus on chromosome 6q25 are associated with breast cancer risk^62^, including in women carrying *BRCA1* and *BRCA2* mutations^63^ and variations in mammographic density^64^, probably through their regulation of *ESR1* gene expression^65^. In breast cancer cells we found enrichment for several notable genes and TFs involved in cancer development including *RB1* and *FOXA1*. The critical role of *RB1* (retinoblastoma protein 1) as a driver of neoplastic development is unequivocal. Alterations of *RB1* or components regulating the CDK-RB-E2F pathway have been identified in nearly every human malignancy and lead to excessive cell growth through loss of cell cycle control. FOXA1 (Forkhead box protein A1) is a member of the forkhead class of DNA-binding proteins and hepatocyte nuclear factor transcriptional activators. It has a well characterised role in breast cancer development acting as a pioneer factor in estrogen receptor (ER) positive breast cancer, where its expression may indicate ER+ cancers that undergo rapid reprogramming of ER signaling are associated with poor outcomes and treatment resistance ^66^. In ovarian cancer cells TFs that were enriched included *MYC, NR2F1 and KDM5B*. Genomic amplification of *MYC* is known to be a major driver of ovarian cancer development^67,68^ while the *CMYC* locus on chromosome 8q24.1 is the most significant pleiotropic risk locus so far identified in breast, ovarian and several other cancers^12,69–73^. *NR2F1* (nuclear receptor subfamily 2 group F, member 1) binds directly with ESR1 and is a member of the nuclear hormone receptor family of steroid hormone receptors, consistent with the underlying etiology and known role of steroid hormones in the risk of ovarian cancer^1,2^. Finally, *KDM5B* (*JARID1B*) regulates *EMSY* via its interaction with the micro-RNA *miR-31*^*74*^ and induces re-expression of *HEXIM1*, which was also enriched in in RCCD1 *trans*-interacting regions in ovarian cancer cells^75^.

Differential gene expression analysis after *RCCD1* overexpression identified 82 genes in both breast and ovarian cancer precursor cell line models that may be associated with the early stages of neoplastic development of breast and ovarian cancers (**Supplementary Table S17**). As expected, *RCCD1* was one of these genes. For others, there is substantial evidence they are altered during breast/ovarian cancer initiation and development including: CCND2 (Cyclin D2), a regulator of cyclin-dependent kinases that appears to be frequently methylated in breast cancers affecting drug resistance^76,77^ and in which common germline variants may affect both risk and survival for ovarian cancer ^78,79^; WNT5A, which is central to many processes in cancer development including cellular senescence, proliferation, invasion, migration, inflammation, metastasis and chemo-resistance^80^, increases cell migration and invasion in breast cancers^81-82^ and is upregulated in ovarian cancer promoting epithelial-mesenchymal transition^83,84^; and the cadherin CDH11 is overexpressed in about 15% of breast cancers and is associated with poor prognosis ^85^. Some genes may represent novel, possibly overlooked candidate biomarkers for screening and/or targets for therapeutic intervention. These include the Ras Protein Activator Like 1 gene, RASAL1 (Ras pathway activation is a critical component of ovarian, breast and many other cancers^86^); the Wilms Tumor interacting protein, WTIP (the Wilms Tumor 1 gene is a highly specific diagnostic and prognostic marker for ovarian cancers)^87^; the kalikrein -related peptidase KLK5 (one of a family of secreted proteins evaluated as screening biomarkers, including KLK3, or prostate specific antigen, which is routinely used to screen for prostate cancer)^88^; and the death associated protein kinase gene DAPK1, a candidate tumor suppressor gene and drug target that promotes the growth of P53 mutant cancers^89^.

The 82 differentially expressed genes shared between breast and ovarian cancer models were also significantly enriched in three of the most well established pathways in cancer: TP53, Hippo and TNF signaling. Aside from TP53, which has been discussed at length already, the Hippo pathway is involved in regulating cell proliferation and promoting apoptosis in organ development and has a critical role in stem cell and tissue specific progenitor cell self-renewal and expansion^90^. Many of the pathway components in the Hippo signaling network are recognized as tumor suppressor genes and oncogenes that are perturbed in human cancers, including Fat4 which is mutated in breast cancer^91^ and YAP in ovarian cancers^92,93^. TNF (tumor necrosis factor) signaling has a primary role in the regulation of immune cells and is involved in a wide range of physiological processes including inducing fever, apoptotic cell death, cachexia, inflammation and tumor inhibition. Dysregulation of TNF production has been implicated in cancer development^94^ with breast and ovarian cancers often accompanied by inflammatory processes characterized by the presence of proinflammatory cytokines including TNF-α^95,96^. Immunotherapy has become a major focus of targeted treatments against a wide range of cancers and within this field members of the TNF receptor superfamily have also been proposed as novel therapeutic targets^97^. Some of the pathways were the same as those identified as enriched networks in *RCCD1* 4C *trans*-interaction studies, in particular steroid hormone biosynthesis in breast cancer and P53 associated pathways in ovarian cancer, even though these networks were derived using different methods and experimental models at different stages of disease pathogenesis.

Strikingly, our data provide evidence that connects 4C *trans*-interaction networks and gene level associations from modelling *RCCD1* overexpression to credible causal risk variants at already identified breast, ovarian and pleiotropic breast-ovarian risk regions. Our results suggest cross-talk between regulatory elements in risk regions with gene promoters at other risk regions on different chromosomes. Recently, Pritchard and colleagues ^98^ proposed a model where genetic contributions to complex traits are partitioned into direct effects from core genes and indirect effects from peripheral genes acting in *trans*. The data we present here indicates that there are likely to be direct functional *trans*-relationships between different GWAS risk loci for any given phenotype. Interestingly, past studies have shown these types of contacts are known to be within interchromosomal enhancer–promoter networks in developmental cell differentiation^99,100^ and have been proposed to contribute to transcriptional heterogeneity within cell populations^101,102^. In one study, aberrant *TAL1* expression in human T-cell acute lymphoblastic leukemia (ALL) was mediated by interchromosomal interaction between the *TAL1* promoter on chromosome 1 with a regulatory element called *TIL16* on chromosome 1^103^ indicating that *trans*-interactions can capture a target gene’s ability to co-regulate the expression of other genes. This supports the hypothesis that weak *trans*-eQTL SNPs contribute to their modest heritability effects through their regulation of peripheral genes to alter core gene expression^98^. If these core genes tend to be co-regulated, then the model also predicts that peripheral variation effects can be increased with a large proportion of heritability driven by these weak *trans* effects.

In summary, we describe a functional framework for the analysis of pleiotropic GWAS risk loci, based on the identification of *trans*-interaction networks combined with differential gene expression modeling of candidate susceptibility genes, to connect different susceptibility loci through their shared biology. The data we present supports the hypothesis that RCCD1 is the susceptibility gene target for both breast and ovarian cancer at the 15q26.1 pleiotropic risk locus, and suggests that by looking at RCCD1 interaction networks throughout the genome we can identify additional risk loci and susceptibility genes for these phenotypes. This may be particularly relevant for studying rare phenotypes where the ability to identify new genome wide risk associations based on case-control genotype analyses alone is restricted by sample size.

## Data Access Statement

**Genome-wide summary genetic association statistics from BCAC are available at:** http://bcac.ccge.medschl.cam.ac.uk/bcacdata/oncoarray/gwas-icogs-and-oncoarray-summary-results/

**Genome-wide summary genetic association statistics from OCAC are available at:** https://www.ebi.ac.uk/gwas/downloads/summary-statistics

## For GTex

The GTEx Project was supported by the Common Fund of the Office of the Director of the National Institutes of Health, and by NCI, NHGRI, NHLBI, NIDA, NIMH, and NINDS. The data used for the analyses described in this manuscript can be obtained from dbGaP via accession number phs000424.

## For TCGA

The results published here are in part based upon data generated by the TCGA Research Network: https://www.cancer.gov/tcga.

## For BCAC

The breast cancer genome-wide association analyses were supported by the Government of Canada through Genome Canada and the Canadian Institutes of Health Research, the ‘Ministère de l’Économie, de la Science et de l’Innovation du Québec’ through Genome Québec and grant PSR-SIIRI-701, The National Institutes of Health (U19 CA148065, X01HG007492), Cancer Research UK (C1287/A10118, C1287/A16563, C1287/A10710) and The European Union (HEALTH-F2-2009-223175 and H2020 633784 and 634935). All studies and funders are listed in Michailidou et al (2017)^16^.

## For OCAC

The ovarian cancer GWAS meta-analyses was particularly supported by US National Institutes of Health grants CA1×01HG007491-01 (C.I. Amos) and U19-CA148112 (T.A. Sellers). The Ovarian cancer Association Consortium has been funded by an Ovarian Cancer Research Fund collaborative grant (A. Berchuck). The COGS project was funded through a European Commission’s Seventh Framework Programme grant (agreement number 223175 - HEALTH-F2–2009-223175). All studies and funders are listed in Phelan et al, 2017^17^.

---

**Supplementary Table S1:** Primers used for luciferase enhancer assay to create risk and non risk SNPs (rs802840, rs8037137)

**Supplementary Table S2:** Colocalization fine-mapping of the breast and ovarian cancer datasets at the 15q26 breast-ovarian cancer risk locus identifies 17 credible SNPs (rows 2-18) explaining > 99% of the shared genetic association in the region.

**Supplementary Table S3:** Quantitative trait locus (QTL) analysis for all 17 credible causal SNPs in the 15q26 risk region and their association with expression of the RCCD1 gene and the PRC-AS1 lncRNA within the region.

**Supplementary Table S4:** Variant effect prediction of altered TFBSs by broken TF motifs of risk SNPs which intersect with 4C interactions using PERFECTOS-APE.

**Supplementary Table S5:** Epigenomic annotations of all 17 credible causal SNPs in the 15q26 breast and ovarian cancer risk region by tissue type. Chromatin state annotations were compiled from H3K27Ac (active promoter/enhancer), H3K4me3(active promoter), H3k4me1 (poised enhancer/promoter), CTCF (architectural protein)

**Supplementary Table S6:** *cis*-4C interaction enrichments in breast, ovarian, and combined breast-ovarian normal and cancer cell lines.

**Supplementary Table S7:** *trans*-4C interaction enrichments in breast (MCF7, BT549) and ovarian (UWB1.289, Kuramochi) cancer cell lines (FDR<0.01).

**Supplementary Table S8:** TFBS enrichment of *trans*-interactions from breast cancer cell lines (MCF7, BT549) in ReMAP 2020 dataset (P<0.01).

**Supplementary Table S9:** TFBS enrichment of *trans*-interactions from ovarian cancer cell lines (UWB1.289, Kuramochi) in ReMAP 2020 dataset (P<0.01).

**Supplementary Table S10:** Differentially expressed genes, ranked by P-value, identified by RNA-Seq analysis, after overexpression of RCCD1 in breast and ovarian precursor cell lines(MCF12A^RCCD1+^, FT282^RCCD1+^ respectively).

**Supplementary Table S11**: KEGG pathway enrichment in breast precursor cells (MCF12A^RCCD1+^)

**Supplementary Table S12:** KEGG pathway enrichment in ovarian precursor cells (FT282^RCCD1+^)

**Supplementary Table S13**: KEGG pathway enrichment in overexpression of RCCD1 in breast (MCF12A^RCCD1^) and ovarian (FT282^RCCD1^) precursor cell lines

**Supplementary Table S14:** Genes with P_adjusted_ < 0.05 and PMAGMA < 2.6 × 10-6

**Supplementary Table S15:** TF Enrichment from LISA Cistrome Analysis in MCF12A^RCCD1+^

**Supplementary Table S16:** TF Enrichment from LISA Cistrome Analysis in FT282^RCCD1+^

**Supplementary Table S17:** Directional effects of DEGs shared between RCCD1 overexpression in breast and ovarian precursor cells (MCF12A^RCCD1+^ and FT282^RCCD1+^); (adjusted P value<0.05)

**Supplementary Figure 1: (A)** RT-PCR analysis of breast and ovarian normal and cancer cell lines. RCCD1 shows higher expression in triple negative breast cancer cell lines compared to normal mammary epithelial cells, in low grade serous ovarian cancer cell lines compared to the likely precursors -fallopian tube secretory epithelial cells and ovarian surface epithelial cells;**(B)** Western blot analysis after CRISPR/Cas9 knockout of RCCD1 in breast (MCF12A^RCCD1-^) and ovarian (FT282^RCCD1-^) cancer precursor cells compared to knockout of the olfactory receptor ORF104A. After RCCD1 knockout and clonal selection, RCCD1 is depleted in MCF12A and FT282 cells, but not in cells after knockout of ORF104A; **(C)(i)** RT-PCR analysis confirming increased RCCD1 expression after stable overexpression of full length RCCD1 and clonal derivation in breast (MCF12A) and ovarian (FT282) cancer precursor cells. (ii) Western blotting analysis confirms over expression of RCCD1 protein in breast (MCF12A^RCCD1+^) and ovarian (FT282^RCCD1+^) cancer precursor cells

