## Supplementary figures and images for "Transcriptome and interactome analyses identify the *TP53* interacting gene *RCCD1* as a candidate susceptibility gene at the 15p26.1 breast and ovarian cancer risk locus"

### Supplementary Figure 1

#### SUPPLEMENTARY FIGURE 1

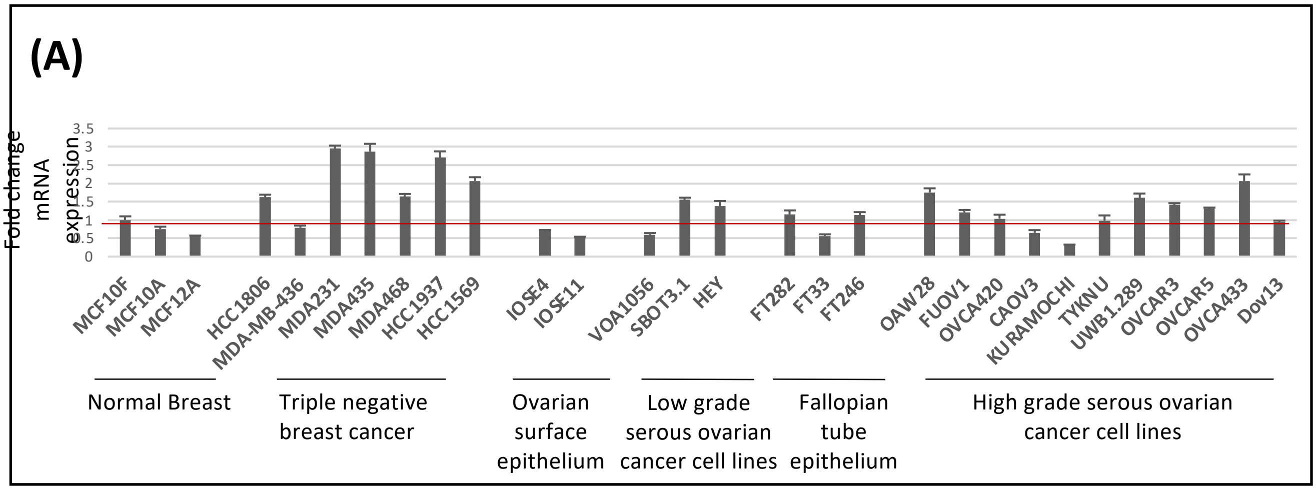


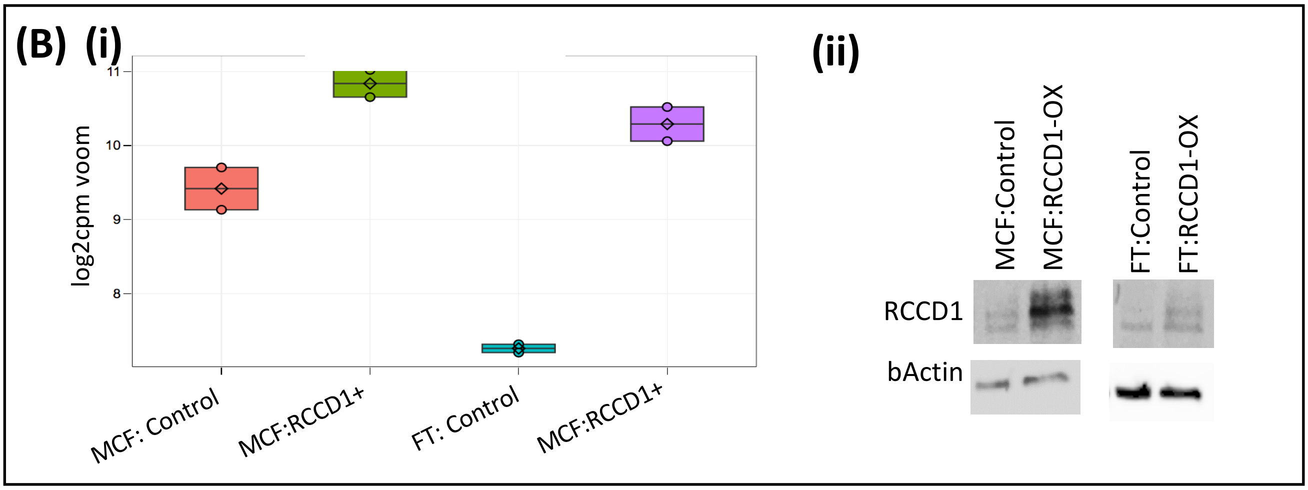


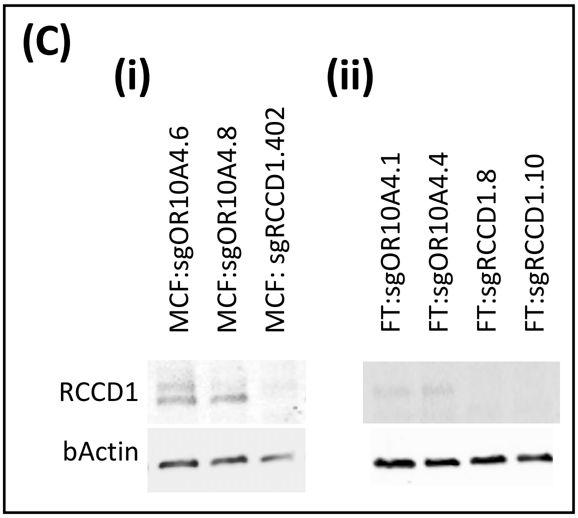
